# CHANGES IN MUSCLE QUALITY FOLLOWING SHORT-TERM RESISTANCE TRAINING IN OLDER ADULTS: A COMPARISON OF ECHO INTENSITY AND TEXTURE ANALYSIS

**DOI:** 10.1101/2024.07.30.605815

**Authors:** Kevan S. Knowles, Jason I. Pagan, Jonathan P. Beausejour, Scott J. Mongold, Abigail W. Anderson, Jeffrey R. Stout, Matt S. Stock

## Abstract

**Background:** Skeletal muscle echo intensity (EI) is associated with functional outcomes in older adults, but resistance training interventions have shown mixed results. Texture analysis has been proposed as a novel approach for assessing muscle quality, as it captures spatial relationships between pixels.

**Purpose:** To examine changes in first-order (EI) and second-order (texture) features of muscle quality following lower-body resistance training in older adults.

**Methods:** Twelve older adults (2 males, 10 females; mean ± SD age = 70 ± 5 years) completed 6 weeks of progressive resistance training, consisting of twice-weekly sessions at 85% of estimated 1RM. Pre- and post-intervention assessments included ultrasound imaging of the rectus femoris (RF) and vastus lateralis (VL), 5-repetition maximum (5RM) leg extension strength, and maximal voluntary isometric contraction (MVIC) force. Ultrasound images were analyzed for EI and texture features using gray-level co-occurrence matrix (GLCM) analysis.

**Results:** Large improvements were observed in 5RM leg extension strength (*p* < 0.001, *d* = 2.09), MVIC force (*p* = 0.006, *d* = 0.969), and RF EI (uncorrected: *p* = 0.003, *d* = 0.727; corrected: *p* = 0.012, *d* = 0.864). No significant changes were observed in muscle size, VL EI, or texture features for either muscle.

**Conclusions:** Short-term resistance training improved strength and RF muscle quality as measured by EI. However, texture analysis features were not sensitive to changes following training. These findings suggest that traditional EI measures may be more appropriate than texture analysis for tracking changes in muscle quality following resistance training in older adults.

**NEW & NOTEWORTHY:** We studied muscle quality changes after 6 weeks of resistance training in older adults, comparing traditional EI to texture analysis. Despite significant strength gains, only RF EI improved, with minimal changes in muscle size, VL EI, or texture parameters. Surprisingly, texture analysis was less sensitive than traditional EI in detecting muscle quality changes. Our results suggest that conventional EI measures may be more suitable for evaluating short-term adaptations to resistance training in older adults.

## INTRODUCTION

The gradual, age-related loss of skeletal muscle mass and strength, a condition known as sarcopenia, will become increasingly pervasive as the global population ages (1, 2). Sarcopenia impairs functional capabilities, leading to adverse outcomes such as falls, fractures, and mortality in older adults (3, 4). The aging process is strongly related to the development of sarcopenia, but lifestyle factors like inadequate nutrition, low physical activity, and poor sleep can accelerate its development (5). While many scientists and clinicians associate sarcopenia with low muscle mass, prospective, longitudinal studies suggest that the loss of muscle strength is ∼3× as rapid as the loss of muscle mass (6, 7). Given this divergent timeline, many have pointed to muscle quality (strength relative to muscle mass) as a more important indicator of functional status among older adults (8). While the exact mechanistic contributors to muscle quality are still under investigation, increases in inter- and intramuscular adiposity have been associated with poor functional outcomes (9) and increased disease risk (10). Given the importance of assessing muscle quality throughout aging, accessible tools that provide insights into these characteristics are of significant value to the scientific community.

Through the use of B-mode ultrasonography, skeletal muscle quality can be non-invasively estimated by quantifying the echo intensity (EI) of a specific region of interest (11). As skeletal muscle consists of both contractile proteins and non-contractile components, (i.e. intramuscular adipose and fibrous tissue) (12), conducting a gray scale analysis of a region of interest can be used to quantify the mean pixel intensity, with black and white pixels corresponding to high and low muscle quality, respectively (13–15). EI of the quadriceps among older adults has been strongly associated with several functional outcomes, including maximal knee extension torque (16), countermovement jump height (17), and normal and fast gait speed (18, 19). While EI has shown promise in cross-sectional studies, investigations that have examined changes following resistance training have shown mixed results. For example, Yoshiko et al. (20) found that after 10 weeks of combined walking and resistance training, there was a significant decrease in EI in the quadriceps femoris among older adults. Radaelli et al. (21) also observed a significant decrease in EI in both upper and lower extremity muscles among apparently healthy adults, regardless of whether high or low volume resistance training was performed. In contrast, Scanlon et al. (22) found that there were no differences in EI for the rectus femoris (RF) and vastus lateralis (VL) in older adults following 6 weeks of progressive resistance training. The reasons why exercise and resistance training studies have produced inconsistent changes in EI could be related to a variety of factors, including different methodological approaches, research participants, and the nature of the intervention (i.e., training volume, intensity, etc.). Establishing reliable protocols that can be used to track muscle quality changes in older adults following resistance training remains a high priority for the field.

Texture analysis of B-mode images offers a novel approach for characterizing skeletal muscle tissue. Texture analysis can be defined as the mathematical characterization of spatial distribution and pixel intensities within a region of interest (23). These features may be used to describe the inherent textural structure of measures of muscle composition (24). First-order features examine individual pixel intensities while ignoring spatial relationships (25), and are evaluated by using the frequency histogram of pixel intensities. This approach provides a count of the number of times a pixel intensity occurs and can be used to calculate mean, standard deviation, skewness, and kurtosis of an ultrasound image (26). While first-order features are easy to implement and interpret, they ignore the spatial relationship between pixels, which can limit their ability to describe underlying texture features of a muscle. Second-order features resolve this limitation by accounting for both pixel intensity and spatial distribution between two pixels using the grey level co-occurrence matrix (GLCM) texture analysis feature. The GLCM is a mathematical feature which describes the joint occurrence of pairs of pixel intensities at specified pixel intensities. From a GLCM, energy, entropy, contrast, homogeneity, dissimilarity, correlation, and variance are common statistical features which can be extracted for characterizing textures (27).

Despite the potential of texture analysis to assess muscle quality, several key gaps in current knowledge persist. The comparative sensitivity of first-order versus second-order features in detecting changes after resistance training interventions remains unclear.

Additionally, the effectiveness of texture analysis in tracking changes in muscle quality in older adults has not been thoroughly investigated. Therefore, the purpose of this study was to examine changes in first- and second-order (GLCM) characteristics of skeletal muscle quality after lower-body resistance training among older adults. To further characterize the influence that resistance training may have on muscle quality measures, we also examined changes in muscle strength and mass. We hypothesize that both the first-order (EI) and second-order (texture) characteristics would show significant improvements after resistance training, and second-order characteristics would demonstrate greater sensitivity to changes in muscle quality compared to traditional EI measures.

## MATERIALS AND METHODS

### Research Design

The data presented herein represents a subset of a larger, more comprehensive investigation (28). Participants completed a supervised, 6-week training program in which they were randomly assigned to either a traditional or functional resistance training group. Participants visited the laboratory for bilateral resistance training twice per week, with visits >48 hours apart. Participants first became acquainted with the exercises and testing protocol during a familiarization visit and engaged in two separate pre-testing visits prior to the intervention. A comprehensive testing battery was used to evaluate pre-post changes, which included ultrasonography of the VL and RF, maximal voluntary isometric contraction (MVIC) force of the quadriceps, and 5 repetition maximum (5RM) strength, among others. The order in which the assessments have been described below corresponds to the order in which they were conducted. Testing sessions were conducted at the same time of day (± one hour) to minimize diurnal effects. Our study design carefully controlled for or monitored hydration levels, nutrition, and physical activity outside of the laboratory. For full details, the reader is directed to the work of Pagan et al. (28).

### Participants

Data from 12 older adults (2 males, 10 females; mean ± SD age = 70 ± 5 years, body mass index [BMI]: 26.5 ± 4.0 kg/m^2^) has been presented in this manuscript. Participants were recruited from the local community with flyers, word of mouth, social media, and posting a study link on the university website. Participants were screened during a phone call to determine eligibility. Exclusion criteria included lower body surgery or use of an assistive walking device within the past year, history of cancer, neuromuscular or metabolic diseases, and myocardial infarction within the past year. Eligible participants had not engaged in lower-body resistance training for at least six months prior to enrollment. All participants signed informed consent documents prior to engaging in study activities. This study was approved by the University of Central Florida Institutional Review Board (#STUDY00004684).

### B-Mode Ultrasonography Image Acquisition and Analysis

A portable B-mode imaging device (GE Logiq e BT12, GE Healthcare, Milwaukee, WI, USA) and a multi-frequency linear-array probe (12 L-RS, 5 – 13 MHz, 38.4-mm field of view, GE Healthcare, Milwaukee, WI, USA) were used to obtain ultrasonography images of the dominant VL and RF (based on kicking preference) to assess muscle size and quality before and after the training intervention. VL images were captured along the lateral aspect of the thigh, at a 50% of the distance between the greater trochanter and the superior border of the patella. Participants were instructed to lie on their non-dominant side of the body with their legs on top of each other, with their dominant knee joint positioned at an angle of 15°, on a treatment table to capture ultrasonography images of the VL. Participants were then instructed to lie down in a supine position on a treatment table and ultrasonography images of the dominant RF were captured. RF images were captured along the midline of the thigh, at 50% of the distance between the anterior inferior iliac spine and the superior border of the patella. Ultrasonography images of both muscles were captured in the transverse plane using the panoramic function. A high-density foam pad was secured to the participant’s thigh to ensure that the probe moved in a straight line along the transverse plane. Three images were taken of each muscle. The same investigator captured all ultrasonography images.

Following data collection, the images were analyzed using ImageJ software (ImageJ, version 1.51, NIH, Bethesda, MD, USA). The polygon function was used to quantify muscle cross-sectional area (CSA) for the VL and RF. Subcutaenous adipose tissue thickness (cm) over each muscle was measured using the straight line function, and corresponded to the mean of left, center, and right values. A texture-based image analysis was employed to investigate the spatial variation in pixel intensity of both muscles. The GLCM characteristics were examined using a plugin macro (Texture Analyzer v0.4) for ImageJ. This plugin requires rectangle-shaped regions of interest for the GLCM analysis, with the goal of incorporating as much muscle as possible without including aponeurosis or bone. The GLCM computation was performed in four directions (0°, 90°, 180° and 270°), and the resultant values were averaged to mitigate the effect of direction (29, 30). Outcome parameters were the angular second moment (ASM), contrast, correlation, inverse different moment (IDM), and entropy. In short, increased ASM, correlation, and IDM values indicate high homogeneity within an image, and increased contrast and entropy indicate high heterogeneity within an image. Increased homogeneity in skeletal muscle typically indicates more dense muscle (less infiltration of connective tissue and adipose; (30). Mathematical formulas for each parameter can be found in Table 1 of Sahinis and Kellis (30). Reported parameters were averaged across the 3 images at each location.

**Table 1.** Test-retest reliability statistics for each of the dependent variables. These statistics were calculated from two pretesting visits prior to the resistance training intervention (*N*=11). Bold font represents a significant change (*p*<.05) from pretest #1 vs. pretest #2. SEMs have been presented in the same absolute unit of the dependent variable. ICC_3,1_, = intraclass correlation coefficient, model 3,1; SEM = standard error of measurement; MD = minimal difference needed to be considered real.

| Dependent Variable | Pretest #1 | Pretest #2 | *P* | *d* | ICC<sub>3,1</sub> | SEM | SEM (%) | MD |
| --- | --- | --- | --- | --- | --- | --- | --- | --- |
| Leg Extension 5RM (kg) | 28.9±13.5 | 32.9±14.2 | **<0.001** | 0.787 | 0.931 | 3.62 | 11.7 | 10.0 |
| MVIC Force (N) | 424.9±151.3 | 430.5±154.1 | 0.405 | 0.139 | 0.965 | 28.5 | 6.7 | 78.5 |
| VL CSA (cm<sup>2</sup>) | 16.5±6.3 | 16.0±5.2 | 0.248 | 0.193 | 0.927 | 1.55 | 9.6 | 4.30 |
| RF CSA (cm<sup>2</sup>) | 4.6±1.7 | 4.3±1.7 | 0.087 | 0.289 | 0.903 | 0.53 | 11.8 | 1.46 |
| VL Uncorrected EI (A.U.) | 107.6±18.9 | 107.1±18.3 | 0.618 | 0.155 | 0.901 | 7.99 | 6.43 | 22.2 |
| RF Uncorrected EI (A.U.) | 107.3±18.0 | 108.2±18.1 | 0.975 | 0.001 | 0.734 | 9.40 | 10.4 | 26.1 |
| VL Subcutaneous Thickness (cm) | 0.731±0.406 | 0.766±0.369 | 0.353 | 0.293 | 0.953 | 0.086 | 11.5 | 0.238 |
| RF Subcutaneous Thickness (cm) | 0.942±0.437 | 0.900±0.408 | 0.373 | 0.281 | 0.963 | 0.084 | 9.0 | 0.231 |
| VL Corrected EI (A.U.) | 152.5±23.3 | 150.7±22.0 | 0.367 | 0.272 | 0.881 | 8.18 | 5.3 | 22.7 |
| RF Corrected EI (A.U.) | 149.1±20.4 | 149.5±18.9 | 0.411 | 0.247 | 0.582 | 9.02 | 7.0 | 25.0 |
| VL ASM | 0.000573±0.000102 | 0.00050±0.000107 | 0.153 | 0.466 | 0.139 | 0.000098 | 18.1 | 0.00027 |
| RF ASM | 0.000645±0.000158 | 0.000698±0.000251 | 0.311 | 0.322 | 0.331 | 0.00017 | 24.8 | 0.00047 |
| VL Contrast | 353.9±125.9 | 367.6±105.6 | 0.607 | 0.160 | 0.772 | 56.6 | 15.7 | 156.9 |
| RF Contrast | 204.4±73.9 | 204.2±86.6 | 0.427 | 0.250 | 0.771 | 36.4 | 18.4 | 101.0 |
| VL Correlation | 0.000573±0.000158 | 0.000535±0.000163 | **<0.001** | 4.47 | 0.431 | 0.00010 | 20.0 | 0.00031 |
| RF Correlation | 0.000775±0.000181 | 0.000872±0.000291 | 0.078 | 0.543 | 0.662 | 0.00016 | 18.8 | 0.00044 |
| VL IDM | 0.160±0.016 | 0.152±0.010 | 0.074 | 0.601 | 0.535 | 0.092 | 5.9 | 0.026 |
| RF IDM | 0.182±0.022 | 0.186±0.027 | 0.312 | 0.321 | 0.668 | 0.014 | 7.4 | 0.038 |
| VL Entropy | 8.06±0.15 | 8.13±0.24 | 0.465 | 0.229 | 0.293 | 1.70 | 2.1 | 0.47 |
| RF Entropy | 7.92±0.22 | 7.86±0.26 | 0.152 | 0.241 | 0.561 | 0.15 | 1.9 | 0.41 |

### Knee Extensor MVIC Force

A custom-built chair with a load cell (Interface, Inc., Force Measurement Solutions, SSMF Fatigue Rated S-type Load Cell, Scottsdale, AZ, USA) was used to assess MVC force of the dominant quadriceps, at hip and knee joint angles of 90°. The back pad of the seat was adjusted to ensure participants were upright and the knee was aligned with the edge of the chair. A seat belt was fastened across the waist to reduce unnecessary movement from the hips and the ankle was secured with a strap to keep the leg stationary. Before maximal testing, participants performed a submaximal warm-up of 3, 10-second contractions at 50% of their perceived maximal force. Following the warm-up protocol, maximal isometric strength of the dominant quadriceps was assessed during 3 MVICs lasting 5 seconds with a 3-minute rest between each attempt. Participants received strong verbal encouragement and were instructed to kick out “as hard and fast as possible.”

### 5RM Leg Extension Testing

To assess lower body muscular strength, 5RM leg extension testing was conducted. The 5RM testing protocol was adapted from Sheppard and Triplett (31) and Stout et al. (32). Participants completed two warmup sets with a light load (∼65% of perceived maximum) for 5-10 repetitions, followed by 3-5 minutes of rest between each set. The load was then increased an additional 4-8 kg (∼10-20%) and participants were instructed to attempt to complete 5 repetitions. Participants that were able to successfully complete their first 5RM attempt with proper technique were given 3-5 minutes rest before performing another 5RM attempt with the load increased an additional 4-8kg. The testing was completed once the participants were able to achieve a true 5RM, with proper form, within 5 attempts. If a participant failed a 5RM attempt, the load from the previous attempt was recorded as the 5RM. Results from the test were used, in part, to determine training loads for participants.

### Resistance Training

Resistance training loads for each exercise were initially set to 85% of the estimated 1RM. Estimated 1RM was based on recommendations from the National Strength and Conditioning Association (33). At the start of each training session, participants completed warm-up sets prior to working sets. Participants completed the following exercises in order: trap-bar deadlift, leg press, leg extension, and leg curl. Each exercise was completed with all repetitions performed to failure for 3 sets at 85% of the estimated 1RM, with 3 minutes of rest between sets and each exercise. Participants were instructed to perform each exercise with a full range of motion and were given the verbal cue “go as quickly as possible” on the concentric action, with eccentric actions being controlled (∼2-3 seconds). To ensure progressive overload, weight was added to each exercise throughout the study, and adjustments to training loads were done on a set-by-set basis. As 85% 1RM corresponds to roughly a 6RM load (31), the goal for each set was for the participants to perform between 3-9 repetitions for each set. For example, if a participant could only perform 2 repetitions for a given set, weight was removed prior to the next set. Conversely, performing 10 repetitions for a given set would result in more weight being added for the next set. This set-by-set approach has proven successful for inducing rapid gain in strength in previous short-term studies (34, 35).

### Statistical Analysis

We first conducted test-retest reliability analyses for each of our dependent variables using the guidelines described by Weir (36). Specifically, for each dependent variable, the ICC (model 3,1) and standard error of measurement ([SEM] reported in both absolute units and as a percentage of the grand mean) were calculated, in addition to paired samples *t*-test and Cohen’s *d* effect sizes. The ICCs were evaluated based on a reliability scale where ICCs < 0.50 indicated “poor” reliability, ICCs of 0.50 – 0.75 indicated “moderate” reliability, ICCs of 0.75 – 0.90 indicated “good” reliability, and ICCs > 0.90 were indicative of “excellent” reliability (37). We also computed the minimal difference needed to be considered real (MD) using equations described by Weir (), which were then used to determine the extent to which a given change was considered meaningful.

For each dependent variable, we conducted paired samples *t*-tests and computed 95% confidence intervals for mean differences and Cohen’s *d* effect sizes. Cohen’s *d* values of 0.20, 0.50, and 0.80 were used to delineate small, medium, and large effect sizes, respectively (38). An alpha level of 0.05 was utilized to determine statistically significant changes. Statistical analyses were performed using JASP 16.0; (University of Amsterdam, Amsterdam, NL, USA). We did not make direct comparisons between the VL and RF muscles, as such comparisons were not relevant to our primary research questions.

## RESULTS

### Reliability

Table 1 shows the results from our test-retest reliability analyses. On the basis of the ICCs, all measures of muscle strength and size, subcutaneous adipose tissue thickness, and VL uncorrected EI showed excellent test-retest reliability (ICC_3,1_, > 0.901). VL corrected EI, RF uncorrected EI, and RF corrected EI showed good-to-moderate reliability (ICC_3,1_, = 0.582 – 0.881). VL and RF contrast showed moderate reliability, with ICCs of 0.772 and 0.771, respectively. Otherwise, most second-order features showed poor-to-moderate reliability (ICC_3,1_, = 0.139 – 0.668). All dependent variables remained statistically unchanged between the two pretesting sessions (*p*>0.05), except for VL correlation and 5RM strength, which likely improved due to task familiarity.

### Changes in Strength, CSA, EI, and Adiposity

Figure 1 shows JASP raincloud plots for 5RM leg extension strength, MVIC force, and VL and RF CSA. Figure 2 shows JASP raincloud plots for uncorrected and corrected VL and RF EI. Moderate-to-large improvements were observed for 5RM leg extension strength (*p* < 0.001, *d* = 2.09), MVIC force (*p* = 0.006, *d* = 0.969), and uncorrected (*p* = 0.003, *d* = 0.727) and corrected RF EI (*p* = 0.012, *d* = 0.864) following training. In contrast to the RF, for the VL, uncorrected EI (*p* = 0.479, *d* = 0.212) and corrected EI (*p* = 0.504, *d* = 0.200) did not improve. Resistance training had only a small-to-moderate influence on CSA for the VL (*p* = 0.209, *d* = 0.385) and RF (*p* = 0.198, *d* = 0.395). There were also no changes in subcutaneous adiposity over the VL (*p* = 0.669, *d* = 0.127) or RF (*p* = 0.251, *d* = 0.350). Table 2 shows mean ± SD data and all statistical results for these outcomes.

**Table 2.**
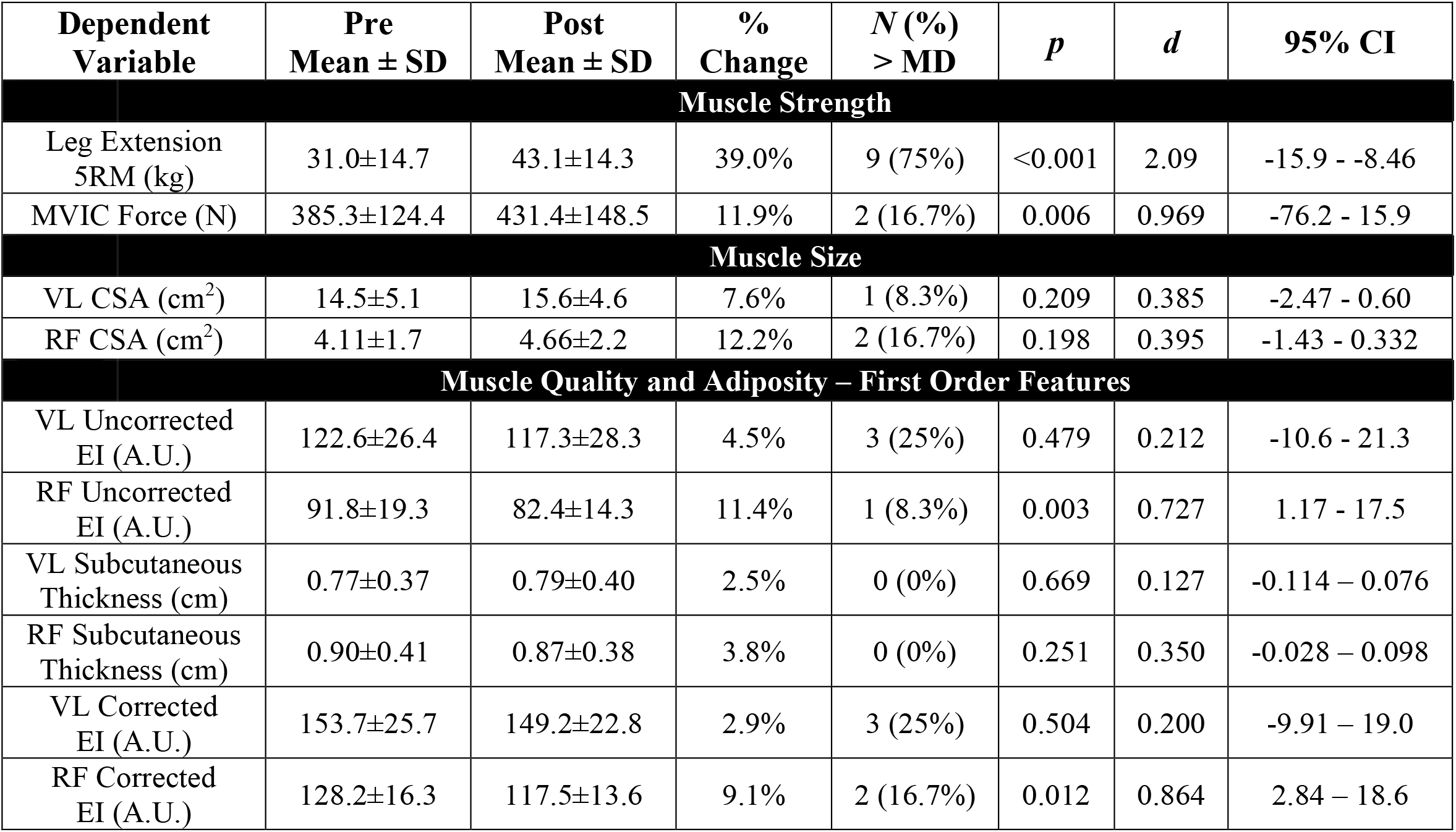
Pretest and posttest data for each of the muscle strength and size and echo intensity (EI) first order features. The fifth column represents the total number and percentage of participants that exceeded the minimal difference needed to be considered real (MD). CI = confidence interval; RM = repetition maximum; MVC = maximal voluntary contraction; RF = rectus femoris; VL = vastus lateralis; CSA = cross-sectional area

**Figure 1.**
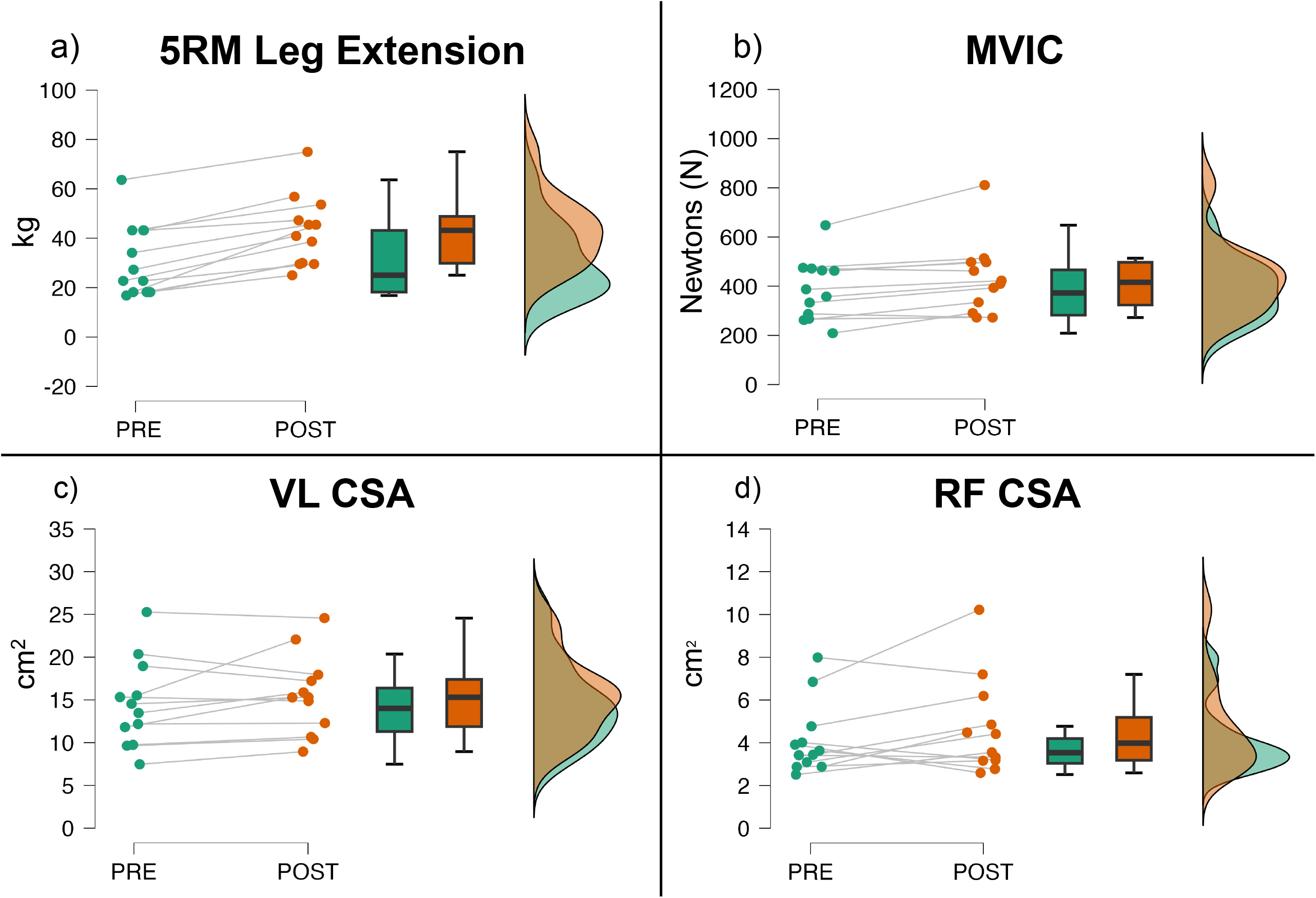
JASP raincloud plots showing pre-post changes in (a.) 5RM leg extension strength, (b.) MVIC force, and CSA for the (c.) VL and (d.) RF following 6 weeks of resistance training in older adults. This plot combines box plots (showing median and interquartile range), density plots (“clouds”), and individual data points (“rain”) with connecting lines. Pre-training values are on the left, post-training on the right. The degree of cloud overlap indicates group changes, while individual points and connecting lines show individual participant responses to the 6-week resistance training program.

**Figure 2.**
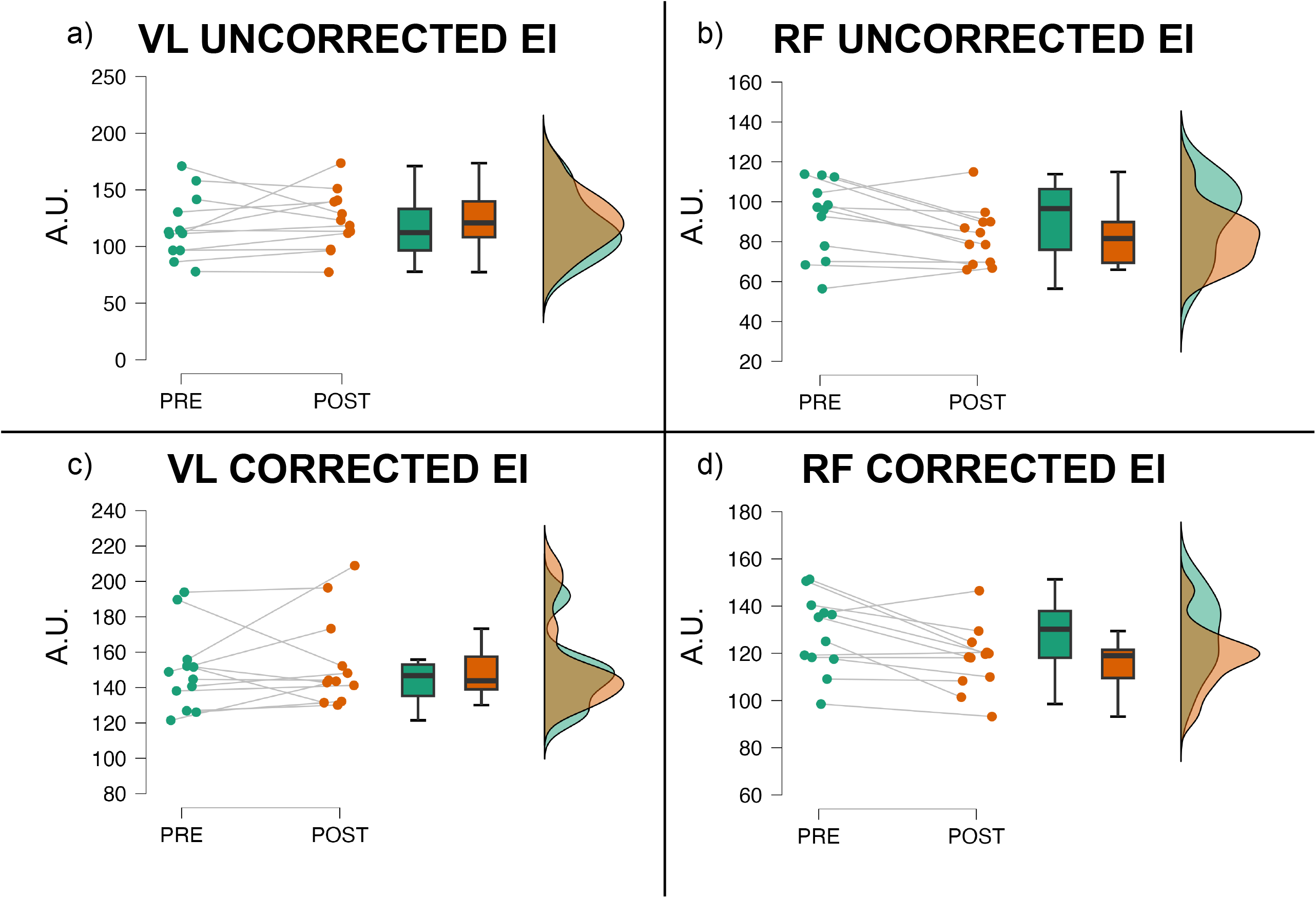
JASP raincloud plots showing pre-post changes in VL uncorrected EI (a.), RF uncorrected EI (b.), VL corrected EI (c.), and RF corrected EI following 6 weeks of resistance training in older adults. This plot combines box plots (showing median and interquartile range), density plots (“clouds”), and individual data points (“rain”) with connecting lines. Pre-training values are on the left, post-training on the right. The degree of cloud overlap indicates group changes, while individual points and connecting lines show individual participant responses to the 6-week resistance training program.

### Changes in Second-Order Features

Figure 3 shows raincloud plots for all second order feature dependent variables for both muscles. None of these variables showed significant improvements following resistance training (*p*>0.05). VL IDM showed a moderate-to-large effect (*d* = 0.737), but all other outcomes showed small changes. The number of older adult participants that showed changes in these variables which exceeded the MD ranged from 0-2. Table 3 shows mean ± SD data and all statistical results for these outcomes.

**Table 3.** Pretest and posttest data for each of the VL and RF second-order features. The fifth column represents the total number and percentage of participants that exceeded the MD. CI = confidence interval; VL = vastus lateralis; RF = rectus femoris; GLCM = gray-level co-occurrence matrix; ASM = angular second moment; IDM = inverse difference moment; MD = minimal difference needed to be considered real

| Dependent Variable | Pre Mean ± SD | Post Mean ± SD | % Change | *N* > MD | *p* | *d* | 95% CI |
| --- | --- | --- | --- | --- | --- | --- | --- |
| **Muscle Quality and Adiposity – GLCM (Second Order Features)** |  |  |  |  |  |  |  |
| VL ASM | 0.0005±0.000107 | 0.000499±0.000101 | 0.2% | 1 (8.3%) | 0.974 | 0.010 | -0.00008149 - -0.00008399 |
| RF ASM | 0.0007±0.00025 | 0.00064±0.0017 | 8.8% | 0 (0%) | 0.187 | 0.406 | -0.00003531 - -0.0001603 |
| VL Contrast | 376.6±105.6 | 354.4±94.6 | 5.9% | 0 (0%) | 0.460 | 0.221 | -24.5 - 50.7 |
| RF Contrast | 204.2±86.6 | 180.5±68.0 | 11.6% | 1 (8.3%) | 0.184 | 0.409 | -13.1 - 60.6 |
| VL Correlation | 0.00054±0.00013 | 0.00052±0.00011 | 3.7% | 0 (0%) | 0.589 | 0.160 | -0.00005623 - 0.00009423 |
| RF Correlation | 0.00087±0.00029 | 0.00080±0.00016 | 8.1% | 2 (16.7%) | 0.430 | 0.237 | -0.000116 - 0.0002536 |
| VL IDM | 0.153±0.010 | 0.157±0.010 | 2.6% | 0 (0%) | 0.207 | 0.737 | -0.008 - -0.0006097 |
| RF IDM | 0.186±0.027 | 0.185±0.023 | 0.5% | 0 (0%) | 0.919 | 0.030 | -0.01 - 0.011 |
| VL Entropy | 8.13±2.44 | 8.17±0.18 | 0.5% | 1 (8.3%) | 0.657 | 0.132 | -0.23 - 0.15 |
| RF Entropy | 7.86±0.26 | 7.94±0.21 | 1.0% | 1 (8.3%) | 0.294 | 0.318 | -0.229 - 0.076 |

**Figure 3.**
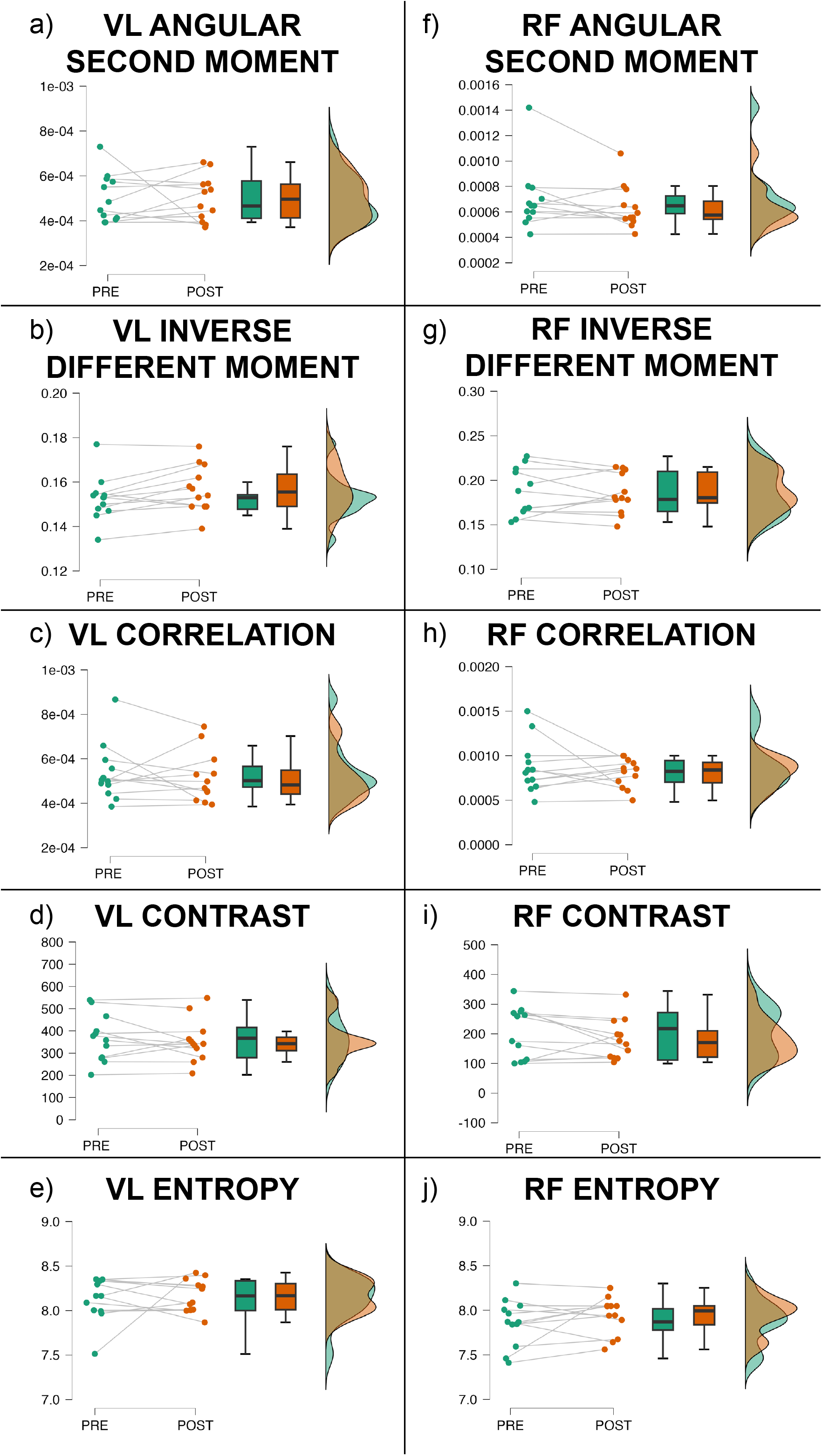
JASP raincloud plots showing pre-post changes in (a.-e.) VL and (f.-j.) RF second-order features following 6 weeks of resistance training in older adults. This plot combines box plots (showing median and interquartile range), density plots (“clouds”), and individual data points (“rain”) with connecting lines. Pre-training values are on the left, post-training on the right. The degree of cloud overlap indicates group changes, while individual points and connecting lines show individual participant responses to the 6-week resistance training program.

## DISCUSSION

Skeletal muscle EI has emerged as an important determinant of functional status among older adults (17, 19). However, resistance training interventions studies exploring changes in EI have shown mixed results, with some reporting significant improvements (20, 21) and others reporting no changes (22, 39). To overcome these inconsistencies, texture analysis has been proposed as a novel, more sensitive approach to assessing EI (24). Therefore, the purpose of this study was to examine changes in first versus second-order texture analysis measures of skeletal muscle quality following progressive resistance training in healthy older adults. As expected, resistance training resulted in substantial improvements in maximal strength, with smaller, non-significant changes in muscle size. Our most important finding, however, was that uncorrected and corrected EI for the RF improved following resistance training, with smaller changes for the VL and no meaningful changes in second-order features. Below we discuss the implications and limitations of our findings, in addition to ideas for advancing research into muscle quality in older adults.

Our data shows a significant decrease in uncorrected and corrected RF EI, suggesting that resistance training may have improved intramuscular adiposity (40, 41), even when accounting for subcutaneous adiposity. These findings are in line with work by Radaelli et al. (21, 42), who found that, after 13 or 20 weeks of progressive lower body resistance training, EI significantly decreased in the RF regardless of whether high or low volume training was performed. Likewise, Yoshiko et al. (20) reported significant decreases in RF EI following 10 weeks of resistance training combined with walking, or just walking alone in older adults. Moreover, Wilhelm et al. (43) observed significant reductions in EI of the quadriceps following 12 weeks of concurrent training. A unique finding of the present study was that the RF showed improvements in muscle quality, whereas the VL did not. These findings are similar to those of Yamada et al. (44) who found that after 12 weeks of progressive bodyweight exercises in community dwelling older adults, RF EI significantly decreased, while the vastus intermedius did not change. This decrease in RF EI, but not the other vasti muscles, may be due to the specific exercises in our training protocol, as the RF may have been activated more due to its role in both knee extension and hip flexion. We believe that the use of both the trap bar deadlift and leg extension may have contributed to the decrease in RF EI, as these exercises may result in greater RF activation due to knee flexion and hip extension (45, 46). Based on these findings, future investigators assessing changes in muscle quality following multi-joint resistance training in older adults should strongly consider examining the RF, in addition to other thigh muscles.

We utilized a novel and comprehensive approach to examine skeletal muscle ultrasound images via texture analysis, which examines the spatial relationship between pixels and intensity of an ultrasound image. However, we observed no significant changes in the chosen GLCM parameters (contrast, correlation, ASM, IDM, entropy) following resistance training. While our work, to the best of our knowledge, is the first to utilize GLCM parameters following a resistance training protocol in older adults, Watanabe et al. (47) conducted a cross-sectional analysis examining differences in GLCM parameters between younger and older adults. They reported significant differences in energy (in this study referred to as ASM) and entropy between age groups, and these features were associated with maximal voluntary isometric knee extensor torque. Specifically, they observed a relationship between a significant decrease in muscle strength and an increase in age and the sum and mean of Haralick features. However, our findings were in opposition to those of Watanabe et al. (47), as we observed a large increase in strength, yet no changes in any of the aforementioned texture features among older adults following resistance training. Yang et al. (48) also found results contradicting our results in their cross-sectional study, where they observed correlations between those with and without dynapenia utilizing EI and texture analysis. They observed that in 36 older adults, texture analysis was able to distinguish muscles of those with and without dynapenia, however, EI was not associated. Based on our findings, we postulate that following a resistance training protocol, EI may be more sensitive to change before any texture analysis parameters. Thus, fine-grained temporal analyses may shed light onto the dynamics of intramuscular adipose, whereby the use of both EI and texture analysis throughout an intervention could clarify whether changes in the amount of adipose precede changes in its spatial organization. While additional research is needed and other findings may be observed under different conditions, our work does not support the notion that second-order features are more sensitive than traditional uncorrected and corrected EI approaches for tracking changes in muscle quality.

Despite its strengths, our study demonstrated strong internal validity and supervised progressive training, several limitations should be considered. This study represents a subgroup of a larger investigation (28), with a small, homogeneous sample of primarily healthy older females, limiting generalizability to clinical populations. The 6-week intervention duration was relatively short compared to studies reporting EI changes after up to 20 weeks of training (20, 21, 42). Although significant strength improvements were observed, longer training periods might yield more pronounced EI changes or eventual changes in textural features. The optimal resistance training paradigms for improving EI in older adults have yet to be established. Our program design, which emphasized high-intensity (85% of estimated 1RM) and low-volume (3-9 repetitions to failure) training, may have prioritized short-term strength gains over whole muscle adaptations. It is possible that manipulating program design variables, such as training volume, intensity, or exercise selection, could be key to improving EI among older adults. To address these limitations and advance our understanding, future research should explore various training protocols to identify the most effective approaches for enhancing muscle quality in this population. Additionally, investigating the potential synergistic effects of resistance training with other interventions, such as protein supplementation or concurrent aerobic exercise, may provide valuable insights. The inclusion of functional performance measures and longitudinal follow-ups could also help elucidate the clinical relevance of EI changes. Furthermore, exploring the relationship between EI improvements and changes in muscle fiber type composition, intramuscular fat content, and connective tissue properties could enhance our understanding of the underlying mechanisms. These considerations should inform the interpretation of our results within the broader context of EI literature and guide future investigations into optimizing resistance training interventions for older adults.

In summary, this study demonstrates that progressive resistance training in older adults leads to significant strength gains and decreases in RF EI, but not in VL EI. Our novel findings reveal that ultrasound texture analysis features are less sensitive to changes following lower body resistance training compared to traditional EI measures in this population. Despite the lack of changes in texture features, resistance training significantly improved lower body strength. These results have practical implications for older adults aiming to enhance lower body strength and reduce fall risk associated with aging. Furthermore, our findings suggest that traditional EI measures may be more appropriate than texture analysis for assessing muscle quality changes following resistance training in older adults. This insight could inform future research methodologies and clinical assessments of muscle quality in aging populations. However, additional research is needed to explore the potential of texture analysis in longer-duration interventions and with different resistance training protocols.

## Notes

**Conflict of interest:** The authors declare no conflicts of interest.

### Competing Interest Statement

The authors have declared no competing interest.

